# Getting more out of FLAG-Tag co-immunoprecipitation mass spectrometry experiments using FAIMS

**DOI:** 10.1101/2021.06.15.448610

**Authors:** Ching-Seng Ang, Joanna Sacharz, Michael G. Leeming, Shuai Nie, Swati Varshney, Nichollas E. Scott, Nicholas A. Williamson

## Abstract

Co-immunoprecipitation of proteins coupled to mass spectrometry is critical for the understanding of protein interaction networks. In instances where a suitable antibody is not available, it is common to graft synthetic tags onto target protein sequences and allowing the use of commercially available antibodies for affinity purification. A common approach is through FLAG-Tag co-immunoprecipitation. To allow the selective elution of protein complexes, competitive displacement using a large molar excess of the tag peptides is often carried out. Yet, this creates downstream challenges for the mass spectrometry analysis due to the presence of large quantities of these peptides. Here, we demonstrate that Field Asymmetric Ion Mobility Spectrometry (FAIMS), a gas phase ion separation device prior to mass spectrometry analysis can be applied to FLAG-Tag co-immunoprecipitation experiment to increase the depth of protein coverage. By excluding these abundant tag peptides, we were able to observe deeper coverage of interacting proteins and as a result, deeper biological insights, without the need for additional sample handling or altering sample preparation protocols.

## Introduction

Many studies in systems biology focus on identifying and characterizing protein-protein interactions (PPI) [1, 2]. Co-immunoprecipitation coupled with mass spectrometry (Co-IP/MS) is an extremely powerful analytical technique and has been pivotal in PPI studies to identify protein complexes, co-factors and signaling molecules [3, 4]. Over the last 20 years, Co-IP/MS approaches have evolved from multiple step isolation procedures such as tandem affinity purification [5] to single step protocols [6] to improve the sensitivity, robustness and ability to assess low affinity PPIs. Despite these improvements, the fundamental approach has remained unchanged and are dependent on the use of specific antibodies against a target protein which exists within a protein complex. Using these affinity reagents, immune complexes are captured and interacting proteins identified using mass spectrometry. While there are thousands of commercially available antibodies and with considerable research efforts being directed toward cataloging these antibodies [7], not all affinity reagents are ideal for Co-IPs or are available for all proteins.

To circumvent the need for protein specific reagents, a common strategy employed by researchers is to tagged proteins with defined peptide or protein sequences. This involves coupling a small peptide or protein tag to the N- or C-terminus of the target protein to minimize its effect on the protein function. The tagged protein can then be detected using antibodies against the tag sequence rather than the target protein itself. These tag-specific antibodies are available commercially and have been very well characterized for their specificity and sensitivity. Examples of common protein tags include green fluorescent protein (GFP), Glutathione S-Transferase (GST) and mCherry while FLAG, c-Myc, 6X-His and Hemagglutinin (HA) are examples of recombinant peptide tags [8–11]. Proteins interacting with the tagged target can be identified by eluting the interacting proteins from the target protein/antibody complex. There are multiple sample preparation options for the elution step and one common approach is to elute the complex from the affinity beads by using competitive displacement [9, 12] whereby molar excess of a synthetic peptide (eg 2X Myc peptide; EQKLISEEDLEQKLISEEDL or 3X FLAG peptide; MDYKDHDGDYKDHDIDYKDDDDK) is added to displace the bound proteins from the tag-specific antibody under non-denaturation conditions. This approach limits the elution of non-specifically bound proteins and allows the elution of complexes in buffers directly compatible with downstream digestion protocols. Unfortunately, the presence of these excess peptides possesses an analytical challenge for subsequent mass spectrometry analysis. Due to the high relative abundances, these ‘contaminant’ peptides rapidly consume the ion storage capacity of ion trapping devices (C-trap, ion trap etc) [13, 14] and thus artificially reduce the ability to detect other peptides (i.e. those arising from interactor proteins of interest) present in the sample mix.

A range of techniques such as gas phased fractionation, BoxCar and Ion mobility (IM) based fractionation [15–18] have been developed that aim to reduce the suppressive effects of highly abundant ions and to increase proteome coverage. Ion mobility mass spectrometry is a technique whereby ions are separated in the gas phase under influence of an electric field. Separation is related to a combination of their size, shape and charge which influence the drift time - the time taken for the molecule to transverse towards the detector [19, 20]. Multiple IM techniques have been developed that differ in the physical principles utilized to achieve ion separation. IM systems can be broadly classified under linear and nonlinear methods [21]. In the linear method, K_0_ (reduced mobility) is assumed to be independent of the electric field. These methods include Travelling Wave (TWIMS), Trapped Ion Mobility (TIMS), Drift Tube Ion Mobility (DTIMS) and Differential Mobility Analyzers (DMA). In the nonlinear methods, K (mobility) of any ion in any gas is dependent on the electric field. These methods include Differential Ion Mobility Spectrometry (DIMS) and Field Asymmetric Ion Mobility (FAIMS). Each IM technique has its strengths and are suitable for different applications. The IM systems can help address some of the issue of under sampling due to the stochastic sampling of eluted peptides and bias caused by poor ion selection in a data dependent acquisition methodology [22]. Multiple groups have utilized different forms of ion mobility mass spectrometry to increase protein coverage in complex cellular systems [16, 18, 23]. We and others have also used IM to enrich for modified peptides [24–26] or as a targeted SRM-like screening [27] technique. These techniques took advantage of the unique collisional cross sectional properties of the peptides on a linear IM device or by matching the applied compensation voltage of the targeted peptide in a nonlinear FAIMS IM device.

FAIMS works by having two electrodes with alternating high and low electric field strength across these electrodes. Separation of ions is by differences in their mobility in high and low electric fields [28, 29]. To prevent collision of the ions with the electrode, the ions are diverted through application of a specific DC compensation voltage (CV). The FAIMS device can therefore be used as a filtering device to remove undesirable singly charged and interfering contaminating ions (eg solvent clusters). This allows the ion current to be spread out and limit the proportion of high abundant species that contribute to the maximal charge capacity of the C-trap [13]. The earlier version of the commercially available FAIMS device has met with limited success due to significant ion losses [30]. Modifications to that FAIMS device by the Moritz lab allowed them to be coupled to nanoLC-MS/MS systems, which significantly increased the depth of proteome coverage in a complex yeast digest [31, 32] . The increased in coverage was attributed to the reduction of singly charged chemical noise and resultant increase in the signal to noise of tryptic peptides. More recent publications from the Coon and Thibaut labs using the second generation FAIMS device [16, 23] upon coupling them to Orbitrap Fusion Tribrid mass spectrometer show marked increase in protein identification from whole cell lysate by ~10-55% than without FAIMS. Similar to the data shown by the Moritz lab, singly charged species are almost constraint to a small CV range of between −10 and −40, are being diverted and are instrumental in the increase in identification.

It is evident that FAIMS has the ability to increase proteome coverage in a highly complex sample by using it as a prefiltering device to remove singly charged chemical noise. The benefit of FAIMS however is not just limited to highly complex samples. In this manuscript, we were able to exploit the same prefiltering principal in a FLAG-Tag Co-IP experiment to filter out highly abundant FLAG peptides that exists in multiple charge states. These highly abundant FLAG peptides would otherwise interfere with the detection of the less abundant peptides derived from the affinity target protein and its respective binding proteins. We were able to demonstrate a huge improvement in terms of identified proteins in a supposedly ‘simple’ Co-IP experiment and the resultant protein-protein interaction network analysis when compared to one without application of FAIMS.

## Experimental Section

### Affinity Enrichment of Flag-Tag proteins

Affinity enrichment of FLAG-Tag protein experiment was performed from eHap knockout cells or HEK293 cells expressing FLAG-Tag protein as described previously [33]. Cell were solubilised in 1% (w/v) digitonin in solubilization buffer (20mM Tris (pH 7.4), 50mM NaCl, 10% (v/v) Glycerol, 0.1 mM EDTA, 5U benzonase (Merck Millipore), 2mM MgCl_2_). Following clarification of cell lysate by centrifugation (20,000*g*, 10 mins, 4°C), 500 μg protein was added to spin columns (Pierce) containing anti-FLAG M2 affinity resin (Merck) and allowed to incubate rotating for two hours at 4°C. Non-specifically bound proteins were then removed by washing with 20x column volumes of ice-cold solubilization buffer containing 0.1% (w/v) digitonin. Proteins were eluted with 50 μl of 100 μg/mL of 3X FLAG tag peptide (Merck) in solubilization buffer followed by acetone precipitation (overnight, −20°C). Proteins were then pelleted by centrifugation (21,000*g*, 10 mins, 4°C), washed with ice-cold acetone and air-dried. The pellet was then resolubilised by sonification in with 8M urea in 50 mM ammonium bicarbonate (ABC). Samples were reduced and alkylated with 50mM TCEP (ThermoFisher) and 500 mM Chloroacetamide (Merck) (30mins, 37°C, shaking). Elutions were diluted to 2M urea in 50mM ABC and digested in trypsin (Promega) overnight at 37°C. The digest was acidified to a final concentration of 1% (v/v) trifluoracetic acid (TFA) and peptides were desalted with stagetips containing 2x plugs of 3M™ Empore™ SDB-XC substrate (SUPELCO). Stagetips were activated with 100% Acetonitrile (ACN) and washed with 0.1% (v/v) TFA, 2% (v/v) ACN prior to binding of peptides. Samples were eluted in 80% ACN, 0.1% TFA and dried completely in a SpeedVac. Peptides were reconstituted in 2% ACN, 0.1% TFA and transferred to autosampler vials for analysis by LC MS/MS

### LC MS/MS analysis

LC MS/MS was carried out using the Fusion Lumos Orbitrap mass spectrometers with the FAIMS Pro interface (Thermo Fisher, USA) and as described previously [26]. The LC system was equipped with an Acclaim Pepmap nano-trap column (Dionex-C18, 100 Å, 75 μm × 2 cm) and an Acclaim Pepmap RSLC analytical column (Dionex-C18, 100 Å, 75 μm × 50 cm). Tryptic peptides were injected into the enrichment column at an isocratic flow of 5 μL/min of 2% (v/v) acetonitrile containing 0.1% (v/v) formic acid for 6 min, applied before the enrichment column was switched in-line with the analytical column. The eluents were 0.1% (v/v) formic acid (solvent A) in water and 100% (v/v) acetonitrile in 0.1% (v/v) formic acid (solvent B). The flow gradient was (i) 0-6 min at 3% B; (ii) 6-35 min, 3-22% B; (iii) 35-40 min, 22-40% B; (iv) 45-50 min, 40-80% B; (v) 50-55 min, 80-80% B; (vi) 55-56 min 85-3% and equilibrated at 3% B for 10 min before injecting the next sample. Tune version 3.3.2782.32 was used. For non-FAIMS experiments, the mass spectrometer was operated in the data-dependent acquisition mode, whereby full MS1 spectra were acquired in a positive mode at 60000 resolution. The ‘top speed’ acquisition mode with 3 s cycle time on the most intense precursor ion was used, whereby ions with charge states of 2 to 7 were selected. Automated gain control (AGC) target was set to standard with auto maximum injection mode. MS/MS analyses were performed by 1.6 *m/z* isolation with the quadrupole, fragmented by CID with collision energy of 35 %, activation time of 10 ms and activation Q of 0.25. Analysis of fragment ions was carried out in the ion trap using the ‘Turbo’ speed scanning mode. Dynamic exclusion was activated for 30 s. For FAIMS-enabled experiments, the mass spectrometer was operated in the data-dependent acquisition mode scanning from *m/z* 300-1600 at 60000 resolution. FAIMS separations were performed with the following settings: inner electrode temperature = 100 °C, outer electrode temperature = 100 °C, FAIMS carrier gas flow = 0 L/min. The FAIMS carrier gas was N_2_. Cycle time using the ‘top speed acquisition’ mode for single CV experiment were 3 s and, for experiments wherin two CVs (−40 and −60) were applied and cycle time was 1.5 s each. MS/MS analyses were performed by 1.6 *m/z* isolation with the quadrupole, fragmented by CID with collision energy of 35 %, activation time of 10 ms and activation Q of 0.25. Analysis of fragment ions was carried out in the ion trap using the ‘Turbo’ speed scanning mode. Dynamic exclusion was activated for 30 s.

For LC MS/MS experiments on an Orbitrap Eclipse the nanoLC conditions were kept constant. The mass spectrometer was operated in the data-dependent acquisition mode, whereby full MS1 spectra were acquired in a positive mode at 60000 resolution. Tune version was 3.3.2782.34. The ‘top speed’ acquisition mode with 3 s cycle time on the most intense precursor ion was used, whereby ions with charge states of 2 to 7 were selected. AGC target was set to standard with auto maximum injection mode. MS/MS analyses were performed by 1.6 *m/z* isolation with the quadrupole, fragmented by CID with collision energy of 35 %, activation time of 35 ms and activation Q of 10. Analysis of fragment ions was carried out in the ion trap using the ‘Turbo’ speed scanning mode. Dynamic exclusion was activated for 30 s.

For LC MS/MS experiments on an Orbitrap Exploris 480, the nanoLC conditions were kept constant. The mass spectrometer was operated in the data-dependent acquisition mode, whereby full MS1 spectra were acquired in a positive mode at 60000 resolution. Tune version was 2.0.182.25. The ‘top speed’ acquisition mode with 3 s cycle time on the most intense precursor ion was used, whereby ions with charge states of 2 to 7 were selected. MS/MS analyses were performed by 1.6 *m/z* isolation with the quadrupole, fragmented by HCD with collision energy of 30%. MS2 resolution was at 15000 Dynamic exclusion was activated for 30 s. AGC target was set to standard with auto maximum injection mode. Dynamic exclusion was activated for 30 s.

### Database search

Database searches was carried out using Proteome Discoverer (v2.4) with the SequestHT search engine or the Maxquant proteomics software package (version 2.0.1.0) against a *Homo Sapiens* database (SwissProt Taxonomy ID 9606, updated Feb 2021). The SequestHT search parameters are Trypsin as the cleavage enzyme and a maximum of 2 missed cleavages. Precursor and fragment mass tolerances of 10 ppm and 0.6 Da, respectively. The default instrument specific search parameters were used in the Maxquant specific searches. For both search engines, carbamidomethyl cysteine was set as fixed modification, and oxidation of methionine and acetylation of the protein N-terminus were considered as variable modifications. Protein and peptides groups were set to a maximum false discovery rate (FDR) of < 0.01 as determined by the Percolator or Maxquant algorithm [34, 35]. Peak area determination was carried out using the Skyline software [36]. Statistical analysis was carried out using the Perseus computational platform [37]. To correct for multiple-hypothesis testing, significant hits are defined using Student’s T-test, truncated by permutation-based FDR threshold of 0.05 (250 randomization) and S0 factor of 2.0 [38]. Protein-protein interaction network and functional enrichment analysis was performed using STRING [39].

## Results

The total ion chromatogram (TIC) of a FLAG Co-IP/MS experiment following competitive displacement with 3X FLAG peptide and *in solution* digest is shown in Figure 1A. Three dominant peaks are visible in the chromatogram between 16.0 min and 26 min. The later TIC peak is being dominated by few highly charged ions, corresponding to multiply charged species of the 3X FLAG peptide - MDYKDHDGDYKDHDIDYKDDDDK (2+, 3+, 4+, 5+ and 6+). The earlier TIC peaks consist of a number of modified or truncated versions of the 3X FLAG peptide e.g. oxidized methionine, trypsin cleavage products (MDYYK↓DHDGDYKDHDIDYKDDDDK) and the oxidized forms of these tryptic peptides. These high abundant peptides typically co-elute with other tryptic peptides within the 10 min elution window. This result is representative of a large fraction of Co-IP/MS experiments wherein the most abundant ions detected are introduced as byproducts of the assay itself and do not correspond to peptides arising from interactor proteins.

**Figure 1.**
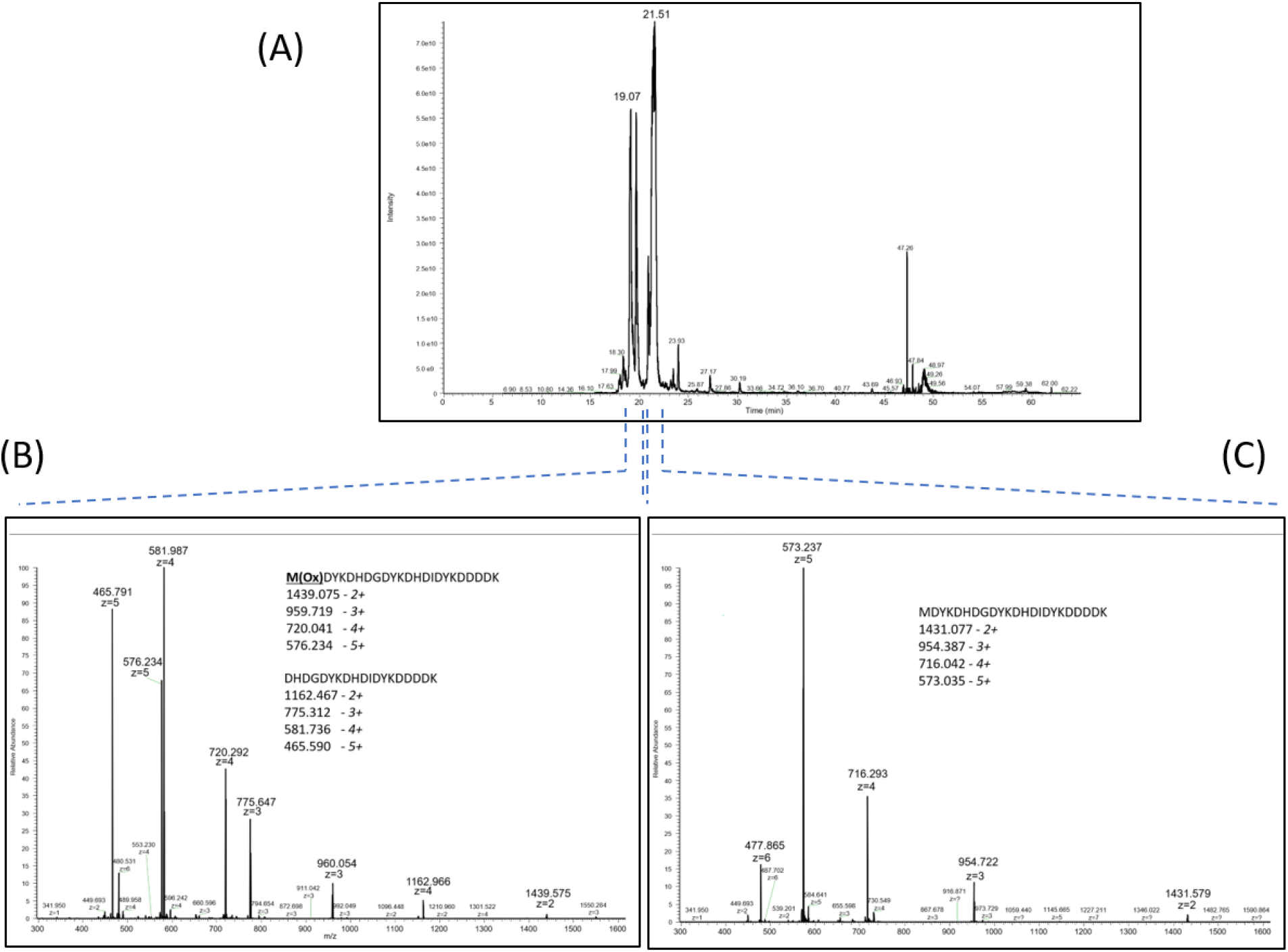
A 3X FLAG tag specific Co-IP Mass spectrometry experiment on an Orbitrap Fusion Lumos mass spectrometer. (A) The TIC mass spectrum showing the the most intense peaks between 18-22minutes. (B) The averaged mass spectrum at ~19min showing the presence of the oxidized version of the 3X FLAG peptide and a trypsin derived truncated version of the 3X FLAG peptide. (C) The averaged mass spectrum at ~21min showing the charge distribution of the intact 3X FLAG peptide. Note m/z peak labeling is on the most abundant ion in the isotopic cluster.

The most abundant ions observed in each of these major chromatographic peaks correspond to highly charged (+4 to +6) 3X FLAG-derived peptide ions (Figures 1B and 1C). Given that tryptic peptides produced from analyte proteins are most commonly observed in +2 or +3 charge states, the higher charges of the 3X FLAG contaminant ions offered the possibility that these could be removed in the gas phase via FAIMS. The synthetic 3X FLAG peptide was first analyzed on the FAIMS/Lumos Orbitrap mass spectrometer with incrementally increasing compensation voltages from CV-20 to CV-70 in steps of 10 (Figure 2). With the exception of the [M+4H]^4+^ precursor at CV-60 and CV-70, all the other charge states of the 3X FLAG were reduced in measured intensity between 30% and 100% (Figure 2, Supplementary Table 1). Next, a FLAG Co-IP experiment was carried out on a whole cell lysate. Following *in solution* digest, the same sample was repeatedly analyzed using the FAIMS/Lumos Orbitrap with the same incrementally increasing compensation voltages to assess the number of proteins, peptides, and peptide spectral matches (PSM) identified under these varying FAIMS conditions. Without FAIMS, 365 proteins, 1362 peptides and 1603 PSM were identified (at 1% FDR). The greatest improvement in identifications were when FAIMS was operated with a CV of −40. This led to gains of 93%, 58% and 51% in proteins, peptides and PSM, respectively, as compared to analysis of the same sample without FAIMS (Figure 3A). The remainder of the single CVs tested led to either comparable (CV- 30 and −50) or substantially lower (CV −20, −60, −70) proteins, peptides and PSM upon database searching.

**Figure 2:**
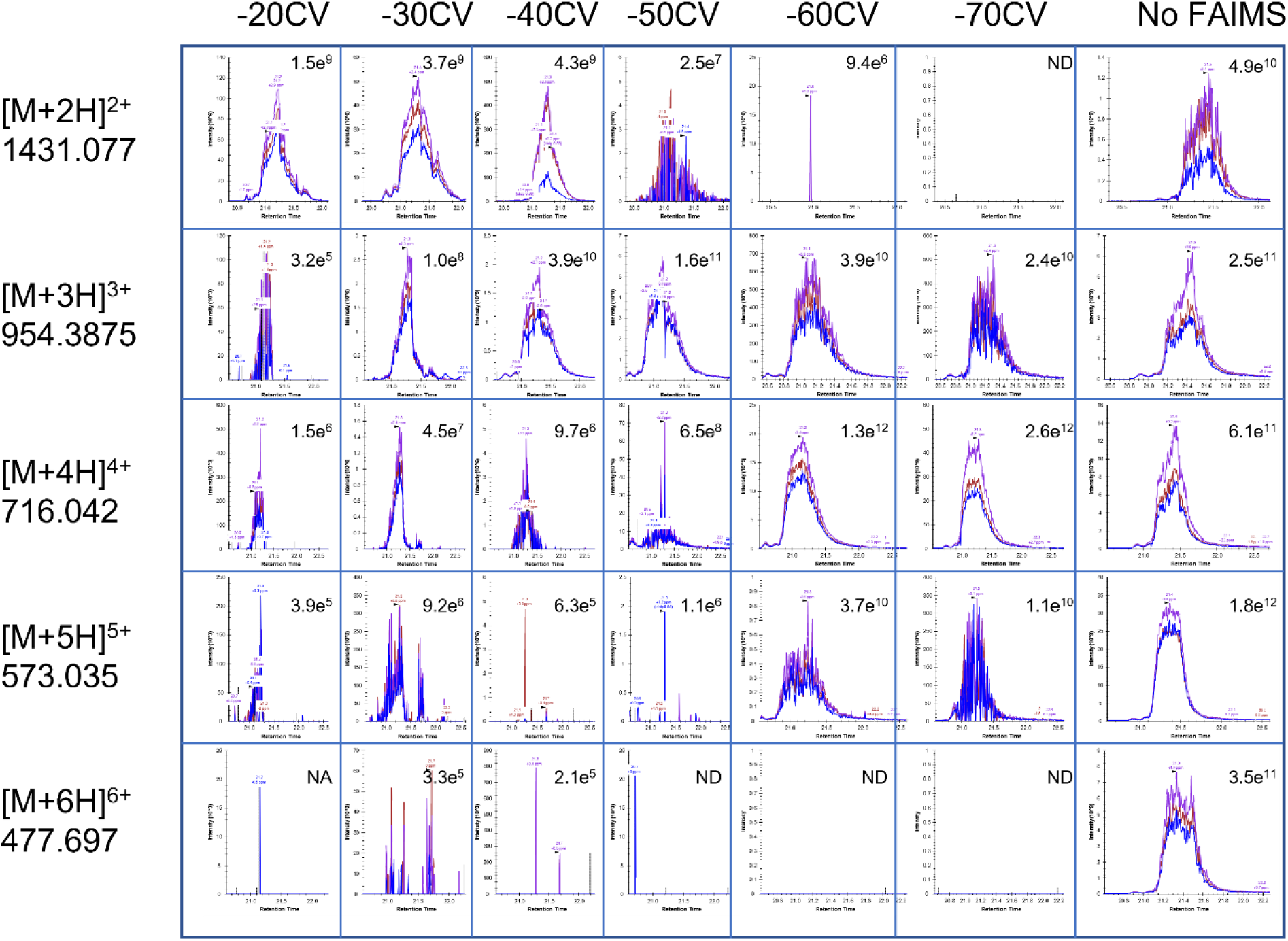
Monitoring the intensities of the different charge species of the 3X FLAG peptide across the different CVs. The extracted ion chromatogram of all graphs is centered at 21.3 min and inserts values represent the total area information. ND = not detected. Metadata is available in Supplementary Table 1.

**Figure 3.**
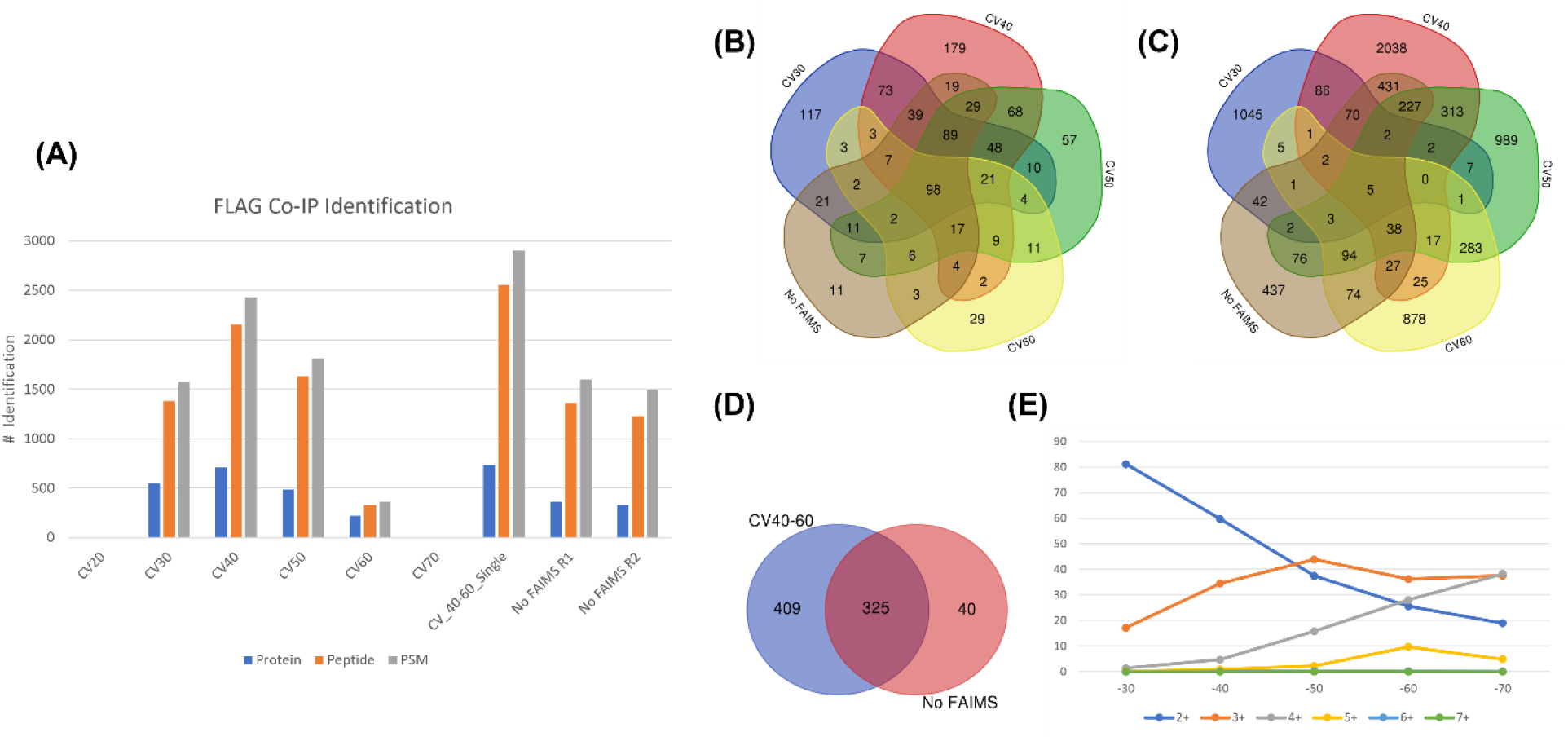
Analyzing the protein and peptide identification from the FLAG Co-IP experiment. (A) Breakdown of protein, peptide and PSM across 6 different CVs, two internal CV stepping and two replicates without FAIMS. We did not identify any proteins with CV-20 and CV-70. (B) Venn diagram of unique protein identification across 4 different CVs (−30, −40, −50 and −60) with No FAIMS applied. (C) Venn diagram of unique peptide identification across 4 different CVs (−30, −40, −50 and −60) with No FAIMS applied. (D) Proteins that are uniquely identified with 2 internal CV stepping vs without FAIMS (E) Charge distribution of MS1 precursors selected for MSMS across the 6 different CVs.

While the greatest improvement in protein identifications was observed for CV-40, it is well documented that different, and perhaps complementary, subsets of peptides may be preferentially detected at alternative CVs or internal CV stepping [23]. To assess whether identification numbers may be improved by employing internal CV stepping, the same FLAG Co-IP digest was re-analyzed with two internal CV steps of −40 and −60. Here, 734 proteins, 2550 peptides and 2904 PSMs were identified following database searching. This translates to a 100%, 59% and 81% gain as compared to without FAIMS. This is almost a 2-fold increase in the total protein identification but critically, the number of proteins that was exclusively identified in the FAIMS experiment was 409 when two internal CV stepping were applied as compared to only 40 without FAIMS (Figure 3D). The distribution of the different charge species selected for MS/MS follows a trend of decreasing 2+ peptides and increasing 4+ and 5+ peptides with more negative CVs (Figure 3E). 3+ peptides appear to be uniformly distributed and within ~20%-45% of all measured MS1 features across CV-30 to CV-70. We repeated this experiment on an independent FLAG Co-IP experiment targeting a different FLAG-Tag protein, and observed similar and large increase in the protein, peptide and PSM (Supplementary Figure 1).

The TIC and number of PSM identifications were plotted across the HPLC retention time at the regions (16-26min) corresponding to the elution time of the 3X FLAG peptides (Figure 1). With FAIMS rapidly switching between CVs of −40 and −60 FAIMS, there is a noticeable drop in the MS1 ion injection time and a corresponding increase in the identified PSM (Figure 4B and 4C). At those alternating CV values, the 5+, 6+ charged species and also a large percentage of the 4+ charged species can be effectively reduced or removed (Figure 2), which otherwise would have contributed to filling up a portion of the limited charge capacity of the C-trap. The total number of identified PSM between the 16-26 minute elution window at CV-40 and CV-60 is 374 and 115, respectively. Without FAIMS, the total number of identified PSM is only 120.

**Figure 4:**
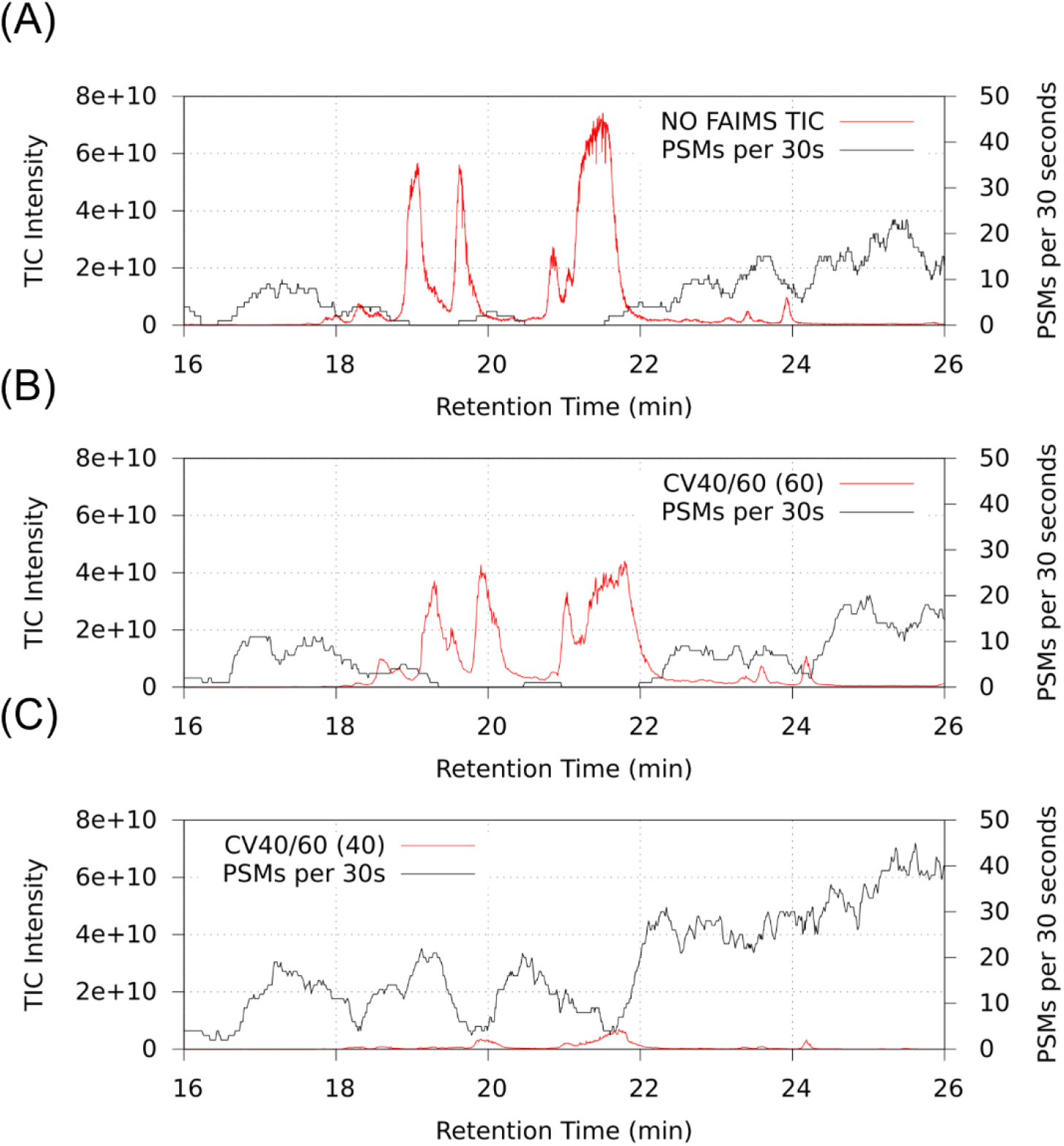
Plot of total ion chromatogram (red trace) and PSM identification (black trace) from the FLAG-Tag Co-IP experiment with (switching between CVs of −40 and −60) and without FAIMS at retention time 16 to 26 min. (A) Experiment without FAIMS (B) Extracted TIC and PSM with FAIMS at CV-60 (C) Extracted TIC and PSM with FAIMS at CV-40.

To determine if sample complexity has a large effect on protein and peptide identification on the Orbitrap Lumos, a commercial Hela tryptic digest (100ng on column) was analyzed using two internal CV stepping (−40 and −60) and without FAIMS. With FAIMS, 37% and 23% more proteins and peptides, respectively, were identified as compared to without FAIMS. To determine if the increase in identification is not due to the analytical mass spectrometer, the FLAG Co-IP digest was analyzed on the next generation Orbitrap Eclipse and Orbitrap Exploris 480 mass spectrometer using similar LC-MS/MS conditions. The results were then compared with the same FLAG Co-IP digest analyzed on the FAIMS/Lumos Orbitrap with two CV internal stepping (−40 and −60). With FAIMS/Lumos Orbitrap, 70.7% and 114.9% more proteins were identified compared to Eclipse and Exploris 480, respectively. On the peptide level, there were 43.3% and 81.1% more from FAIMS/Lumos compared to Eclipse and Exploris 480, respectively (Supplementary Table 2).

All the above Co-IP experiment were carried using the same *in solution* digested sample and comparing protein and peptide identifications, with and without FAIMS. We sought to see if the improvement were indeed biologically relevant in a comparative Co-IP experiment. For this, a FLAG-Tag transmembrane protein was used for immunoprecipitation of proteins after treatment with a drug that targets the membrane ion channel. All experiment were performed in triplicates and the *in solution* digested samples were analyzed on the Lumos Orbitrap mass spectrometer with and without FAIMS. After subtracting proteins identified from the negative controls, statistical analysis was carried out to highlight proteins that were significantly changed in abundances upon drug treatment (Figures 5A and 5B). These significantly changed proteins were combined with proteins that are identified exclusively from the drug treatment group (identified in all 3 replicates in drug treatment and absent in all the non-treatment replicates). From the above selection criteria, a list of 265 proteins with FAIMS and 117 proteins without FAIMS was generated (Supplementary Table 3). Protein-protein interaction network and functional enrichment analysis was then carried using STRING (Figures 5D and 5E). We applied the highest confidence score (0.900) and display only edges from where proteins are part of a physical complex (STRING physical subnetworks feature). From these, a highly significant protein-protein interaction (PPI) enrichment for both Co-IP with FAIMS (PPI enrichment p value <1.0e-16) and without FAIMS (PPI enrichment p value = 1.3e-11) was obtained. For clarity, the disconnected nodes in the network were not displayed. There are a total of 51 protein (86 edges) enriched from known physical complexes with FAIMS versus 18 proteins (17 edges) from without FAIMS. The top 2 molecular function enrichment from both FAIMS and without FAIMS were related to SNAP receptor (red bubble) and SNARE binding (blue bubble). The endomembrane system (green bubble) was the topmost enriched subcellular localization for both FAIMS and without FAIMS Co-IP experiment.

**Figure 5:**
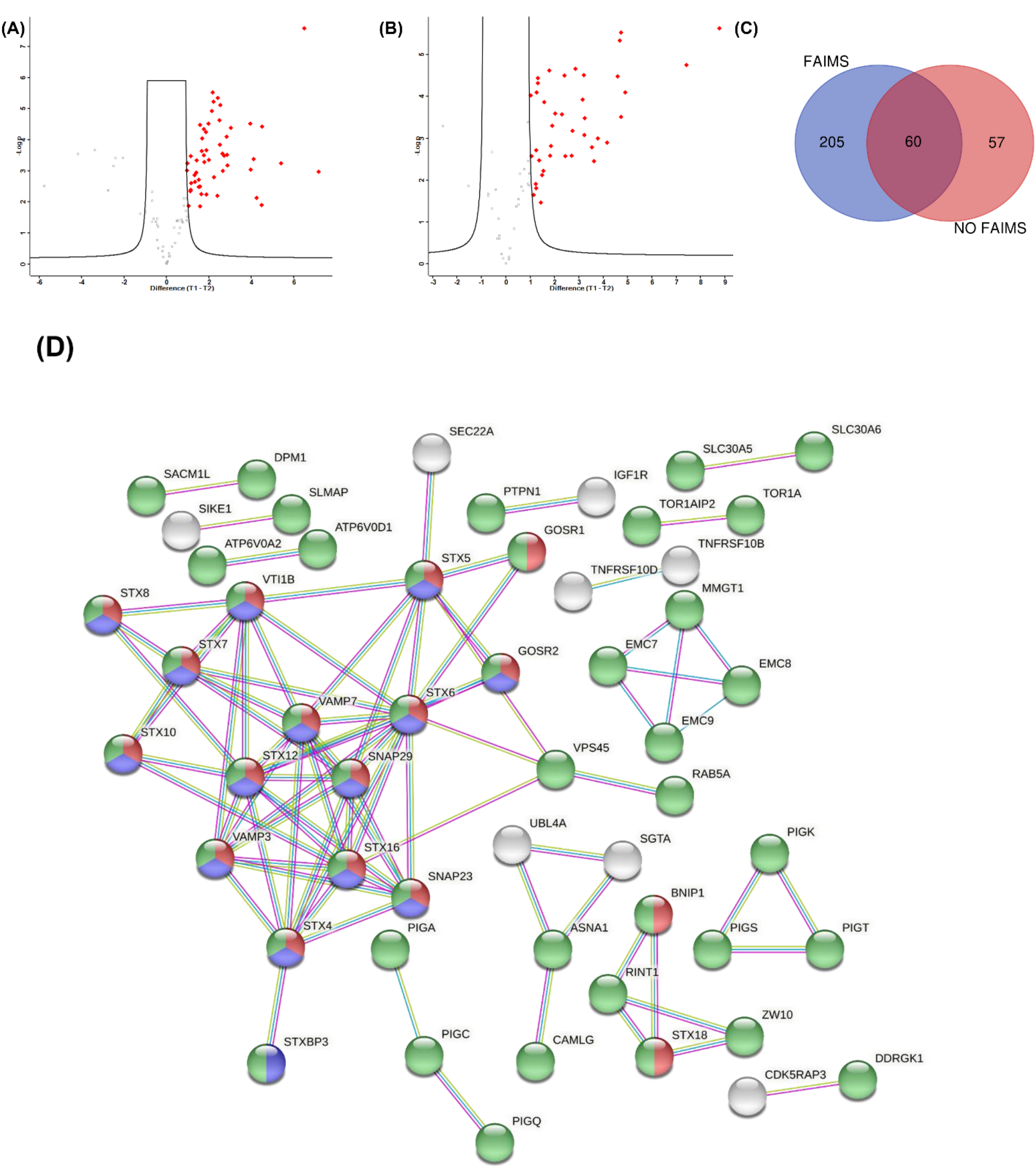
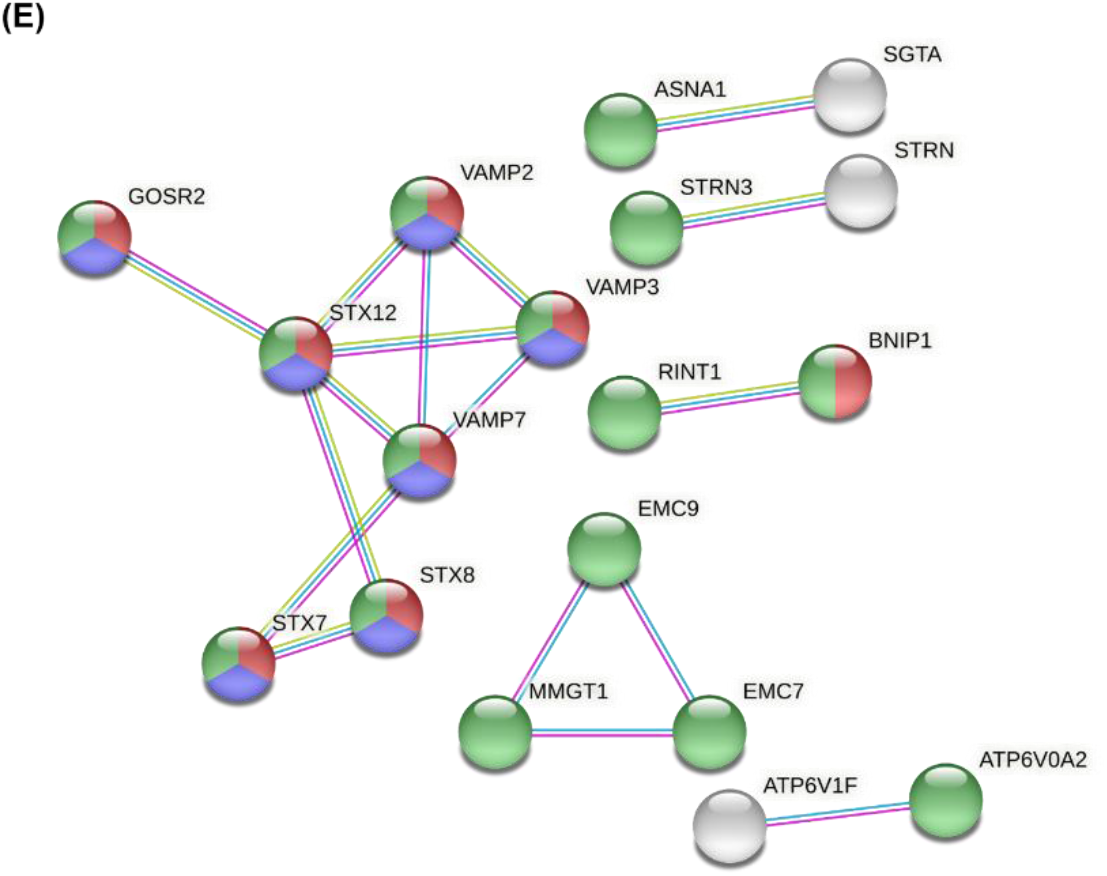
Protein-protein interaction from Co-IP with and without FAIMS (A) Volcano plot from the FAIMS enabled experiment and highlighting proteins (red diamond) that have significantly increased in abundances in drug treated samples (proteins identified in 3 replicates, T-Test, FDR 0.05 and S0=2). X-axis represents log2 transformed fold change and y-axis represents the T-test p values. (B) Volcano plot from the without FAIMS experiment and highlighting proteins (red diamond) that have significantly increased in abundances in drug treated samples (C) Venn diagram of proteins submitted to STRING analysis. These include proteins significantly increased and proteins exclusively found in the drug treatment groups (identified in all 3 replicates of the drug treated group and absent in the other group). (D) STRING physical subnetwork analysis for the FAIMS enabled experiment. For clarity, disconnected nodes have been hidden. Red and blue bubble represent molecular functions enriched in SNAP receptor (p-value = 5.14e-16) and SNARE binding activity (p-value = 3.56e-08), respectively. Green bubble represents enrichment in the endomembrane system (p-value = 1.53e-61). (E) STRING physical subnetwork analysis for without FAIMS experiment. For clarity, discontinued nodes have been hidden. Red and blue bubble represent molecular functions enriched in SNAP receptor (p-value = 2.11e-08) and SNARE binding activity, respectively (p-value = 0.0015), respectively. Green bubble represents enrichment in the endomembrane system (p-value = 1.73e-31). Edges color legend: Pink = experimentally defined, Magenta = from curated databases and Light Green = text mining

## Discussion

Given that FLAG-Tag Co-IP experiments typically produce large molar excesses of contaminant peptides derived from peptides used in competitive displacement, we reasoned that deeper profiling of interacting species could be achieved by developing methods to reduce the suppressive effects of high-abundance ions on mass spectrometric detection of low-abundance species. We have utilized FAIMS as a gas phase filtering technique, to filter or reduce the highly abundant synthetic 3X FLAG peptides in a FLAG Co-IP mass spectrometry experiment and have increased the depth of proteome coverage. The total numbers of proteins identified from FLAG-Tag Co-IPs increased two-fold when FAIMS filtering was employed with two, rapid compensation voltage steps (−40 and −60) compared to analyses of the same sample when FAIMS was not used. Critically, a much larger number of proteins were exclusively identified with the application of the optimal FAIMS-filtering method versus when no FAIMS is used.

It should be noted that a range of techniques are currently available to reduce interference from high-abundance peptides. For example, the excess FLAG peptides could be removed with an offline cleanup process or by running it off an SDS PAGE gel, size exclusion columns, molecular weight filtration etc. The downside is that it requires additional offline sample handling steps and will likely introduce experimental variabilities. A single LC MS/MS experiment with increased depth is still the preferred methodology in a Co-IP/MS experiment. In contrast to these existing methods, the FAIMS methodology presented here does not require any additional sample handling steps, fractionation or library construction and is completely amenable to standard, single-shot data-dependent acquisition.

We reasoned that one of the largest contributing factors to the increased protein identification is that FAIMS can divert singly charged chemical noise, resulting in increased signal to noise as also demonstrated in earlier studies [16, 23, 31, 32]. Not allowing a few abundant peptides, such as the synthetic 3X FLAG peptide fill up the limited charge capacity of the ion storage device is another significant contributing factor. The later phenomenon has been exploited to increase the proteome coverage in other gas phase fractionation strategies [15, 17, 40]. For example, in the more recent BoxCar acquisition strategy [17], ion injection is limited to a portion of the total mass range so as to distribute the maximal charge capacity. This is to limit the proportion of highly abundant or well ionizing peptides affecting ion accumulation in the C-traps, which typically can be filled up in less than 1ms [13]. One millisecond is typically less than 1% of the transient time required to generate a high-resolution mass spectrum in the Orbitrap, which takes between 128-256ms (resolution dependent). What that means is >99% of these ions are not being used for mass analysis. When scanning in limited mass ranges, these gas phased fractionation methodologies can spread out the abundant species which otherwise rapidly fill up the C-trap, thus allowing increased filling time for the less abundant peptides. Although the MS1 overhead and duty is increased with multiple m/z windows, the overall benefits in terms of increased identification of low-level peptides is substantial with up to a 10-fold gain in dynamic range [17]. In a FLAG Co-IP experiment, the presence of the high abundant synthetic 3X FLAG peptide will rapidly fill up the C-trap and can affect accumulation of other low abundant peptides. We have empirically determined the relationship between the applied CV and the charged characteristic for the 3X FLAG peptide (Figure 2). By applying different FAIMS CVs, gas phased fractionation can be used to filter out the high abundant 3X FLAG peptides at different charge states. The charge distribution and mass to charge ratio (m/z) of precursor peptides across multiple CVs in a complex Hela and yeast digest have previously been reported [23, 31, 32]. In addition to the charge states, precursor mass to charge ratio (m/z) also have a large impact on the FAIMS mobility. Optimal compensation field (*Ec*) for multiply charged peptides varies and the largest m/z are generally observed over lower magnitude *Ec* and smaller m/z generally over higher *Ec* and are charge state dependent [32]. The reported Co-IP data is in general agreement with the complex digest reported in the earlier investigations, with doubly charged precursors found predominantly at the lower CV, triple charged precursors distributed over a wider CV and higher charge densities observed for more negative CVs.

When performing that as a single LC-MS/MS experiment and two internal CV stepping, a large proportion of the intense 4+, 5+ and 6+ FLAG peptide is filtered at the CV-40 step. This allows gas phase enrichment of the lower charged state peptides derived from the affinity target protein and its respective binding proteins. When switching to the CV-60 step, the characteristics of the peptides differs, allowing identification of a different set of peptides. This is also evident by the decreased MS1 TIC and corresponding increased PSM during the 16-26 minute elution window of the synthetic 3X FLAG peptide (Figure 4). In contrast to the BoxCar acquisition strategy, the minimal increased in MS1 overhead in a FAIMS type experiment has lower impact on the overall instrument duty cycle as only 2 internal CV stepping are used, and the complexity of Co-IP experiment is generally not high.

We sought to establish if the large increase in protein and peptide identifications is a phenomenon that is augmented in Co-IP experiment and not due to sample complexity or instrumentation. To do that, the differences in identifications using a highly complex whole cell digest with and without FAIMS was carried out. In a complex whole cell digest and using FAIMS, there were 37% and 23% more proteins and peptides identified, respectively, than without FAIMS (Supplementary Table 2). This is in general agreement with the work published by the Coon lab [23] where they showed FAIMS (different CVs compared to this manuscript) providing similar increase in protein identification with the same 60 min analysis time. In the FAIMS Flag-Tag Co-IP experiment, where sample complexity is much lower, there is a 100% and 59% increase in proteins and peptides, respectively. This suggests, removal of singly charged chemical noise and high abundant contaminant 3X FLAG peptides can have a more profound effect in samples of lesser complexity. The same Co-IP experiment was then analyzed without FAIMS on the Orbitrap Lumos and 2 newer Orbitraps (Orbitrap Eclipse and Exploris 480) to determine if the same increase in protein and peptide identifications can be achieved with newer instrumentation. The number of identified proteins, peptides and PSM was similar between Orbitrap Eclipse and Lumos and Exploris 480 having the least number of identifications (supplementary Table 2). Again, when comparing all 3 experiments with FAIMS/Lumos, protein, peptide and PSM identification are much higher when FAIMS was used. The Orbitrap Eclipse and Exploris 480 are to date, the latest generation Orbitrap mass spectrometer with improvement in the quad filter, FTMS overheads and better ion transmission [41, 42]. These newer generation of Orbitrap mass spectrometers have been shown to increase the identification rates of proteins as compared to the Fusion Lumos Orbitrap by up to 20% in a complex Hela digest or single cell proteome study where sensitivity is of upmost importance [41, 43]. The much higher protein and peptide identifications clearly shows the increase identification on FAIMS/Lumos cannot be replicated with newer instrumentation. The FAIMS device can provide increased depth in a highly complex sample but for a relatively low complexity sample like a Co-IP experiment, the improvement afforded by filtering out the dominating ion species is much greater.

Lastly, we look into the STRING protein-protein interaction networks generated from identified proteins with and without FAIMS. It quickly became apparent that the network generated from the FAIMS dataset is denser, more informative and has more proteins identified than without FAIMS (Figure 5C). Whilst not focusing on any biological interpretation, we sought to find out what gains were achieved. Instead of presenting indirect (functional) interactions where the PPI network can become overly dense and with boundless overlapping nodes, these networks are reported as a more conservative physical subnetwork. This allow us to look at proteins that are experimentally shown to be physically linked and without the added complications of functional interactions derived from computational means [44]. For added clarity, disconnected nodes are also removed from the analysis. With FAIMS, there are 51 proteins that are connected through direct physical interactions versus 17 without FAIMS. The vast majority of these proteins are associated with the endomembrane system, which correlates well with the bait protein being a transmembrane protein. The top 2 enriched molecular function from the FAIMS and no FAIMS dataset are similar and are related to SNARE binding and SNAP receptor activity. SNARE (or <u>SNA</u>P <u>RE</u>ceptor) proteins are made up of a large protein superfamily of more than 60 members in mammalian cells and are involved in membrane fusion along the secretory and endomembrane systems [45]. Although analysis without FAIMS showed similar enrichment in molecular functions, more proteins from the SNARE complexes such as Syntaxins (STX), STXBP, VTI1B, VAMP, SNAP, SEC22A, GOSR2 and RAB [46] are enriched with FAIMS. These SNARE proteins are on top of many more interacting proteins, especially those associated with the endomembrane systems (Figure 4D, green bubble). Being able to identify different STX proteins can provide us with additional information on the cellular localization of these complexes. For example, different members of the SNARE proteins are distributed in distinct subcellular localization, which form specific SNARE complexes to mediate different transport events [46]. 9 different STX proteins versus 3 STX proteins were identified from FAIMS and without FAIMS, respectively. STX5 is present in at least three different SNAREpins, regulating Golgi trafficking and is thought to be the master SNARE of the Golgi apparatus [47]. STX6 and STX16 form one of two Golgi SNAREpins with VTILA and VAMP4 [48]. Consolidating the above information allow us to speculate an enhanced interaction of the Golgi network with the transmembrane protein, upon treatment with the membrane ion channel drug. That cannot be said with the Co-IP experiment analyzed without FAIMS. We cannot discount there are also proteins that are uniquely identified from the without FAIMS experiment, but the large increase in protein identification and resultant denser PPI network represents the extension of the depth of proteome coverage, potentially revealing previously unknown protein partners and in our view should be the method of choice for such experiment.

Discovery and functional characterization of protein-protein interaction network is challenging. This requires sound experimental planning, the right bait protein(s), high yield of bait proteins and in-depth identification of the interacting proteins. While a full biological analysis of these entities is beyond the scope of this manuscript, these proteins differentially detected with FAIMS could potentially be missing links in protein complexes or low level transient interactors etc. While FAIMS enabled methodology has already been documented to increase proteome coverage for complex proteomics samples we show here that it can have a proportionately greater impact on “simple” or low complexity samples.

## Data Availability

The mass spectrometry proteomics data have been deposited to the ProteomeXchange Consortium via the PRIDE partner repository with the dataset identifier PXD028965

## Author Contributions

CSA and NAW designed the research, CSA developed methodology, CSA, MGL, SN, SV performed experiments, all authors analyzed data and wrote the paper.

## Supporting information

Supplementary data to this article can be found online at …

**Supplementary Figure 1:** Analyzing the protein and peptide identification from a second independent FLAG Co-IP experiment.

**Supplementary Table 1:** Comparing intensities of the different charge states of the 3X FLAG peptide across different CVs.

**Supplementary Table 2:** Comparing the identification rate across 3 different Orbitrap Mass Spectrometers

Supplementary Table 3: List of significantly changed proteins used for STRING analysis

